# Early evoked brain activity underlies auditory and audiovisual speech recognition deficits in schizophrenia

**DOI:** 10.1101/2021.10.09.463763

**Authors:** Daniel Senkowski, James K. Moran

## Abstract

**Objectives:** People with Schizophrenia (SZ) show deficits in auditory and audiovisual speech recognition. It is possible that these deficits are related to aberrant early sensory processing, combined with an impaired ability to utilize visual cues to improve speech recognition. In this electroencephalography study we tested this by having SZ and healthy controls (HC) identify different unisensory auditory and bisensory audiovisual syllables at different auditory noise levels.

**Methods:** SZ (N = 24) and HC (N = 21) identified one of three different syllables (/da/, /ga/, /ta/) at three different noise levels (no, low, high). Half the trials were unisensory auditory and the other half provided additional visual input of moving lips. Task-evoked mediofrontal N1 and P2 brain potentials triggered to the onset of the auditory syllables were derived and related to behavioral performance.

**Results:** In comparison to HC, SZ showed speech recognition deficits for unisensory and bisensory stimuli. These deficits were primarily found in the no noise condition. Paralleling these observations, reduced N1 amplitudes to unisensory and bisensory stimuli in SZ were found in the no noise condition. In HC the N1 amplitudes were positively related to the speech recognition performance, whereas no such relationships were found in SZ. Moreover, no group differences in multisensory speech recognition benefits and N1 suppression effects for bisensory stimuli were observed.

**Conclusion:** Our study shows that reduced N1 amplitudes relate to auditory and audiovisual speech processing deficits in SZ. The findings that the amplitude effects were confined to salient speech stimuli and the attenuated relationship with behavioral performance, compared to HC, indicates a diminished decoding of the auditory speech signals in SZs. Our study also revealed intact multisensory benefits in SZs, which indicates that the observed auditory and audiovisual speech recognition deficits were primarily related to aberrant auditory speech processing.

**Highlights:** Speech processing deficits in schizophrenia related to reduced N1 amplitudes Audiovisual suppression effect in N1 preserved in schizophrenia Schizophrenia showed weakened P2 components in specifically audiovisual processing

## 1 Introduction

Many patients with schizophrenia (SZ) show impaired language functions, such as disorganized speech (de Boer et al., 2020; DeLisi, 2001). Furthermore, electrophysiological and functional neuroimaging studies in SZ have found aberrant processing of speech at the neural level (Hirano et al., 2020; Li et al., 2007). The vast majority of studies, however, examined unisensory speech processing using, e.g., articulated words or written sentences as stimuli (Javitt and Sweet, 2015; Mohr et al., 2000). In everyday life, speech usually occurs in a multisensory context – viewing lip movements and hearing words. Thus far, studies investigating audiovisual (AV) speech processing in SZ (De Gelder et al., 2002; Roa Romero et al., 2016a, 2016b; Ross et al., 2007b; Stekelenburg et al., 2013; Szycik et al., 2013, 2009) have found evidence for impaired bisensory audiovisual speech processing in SZ. However, the neural basis of these deficits, especially the temporal dynamics, are not well understood. While previous electrophysiological studies have primarily focused on alterations in late stimulus processing (Mathalon et al., 2002; Mohammad and DeLisi, 2013), much less is known about the role of early neural processing for speech recognition deficits in SZ.

Previous research in healthy individuals has shown relationships between the amplitudes of the early auditory evoked N1 and P2 components in event-related potentials (ERPs) and bisensory audiovisual speech processing (Baart et al., 2014; Besle et al., 2008, 2004; Jaaskelainen et al., 2004; van Wassenhove et al., 2005). Analyzing data across various electroencephalography (EEG) experiments, Baart (2016) found a suppression of the N1 and P2 components for bisensory audiovisual compared to unisensory auditory speech stimuli. In particular the suppression of the N1 component has been interpreted as an early multisensory facilitation of auditory speech processing (Brunellière et al., 2013; van Wassenhove et al., 2005). In SZ only one study has thus far examined audiovisual speech integration in early ERPs (Stekelenburg et al., 2013). The central finding of this study is the lack of an audiovisual N1 suppression effect in SZs. The study included a simple visual catch trial task to ensure that participant attended to the visual inputs, but no active speech processing was required. Therefore, it is possible that differences in attention allocated to the speech stimuli contributed to the observed group differences in N1 suppression effects. Moreover, there is not enough information on behavioral data to determine whether the lack of audiovisual N1 suppression in SZs was related to possible speech recognition deficits. To reveal a more accurate picture on the role of early neural processing for auditory and audiovisual speech perception SZ, an experiment with active speech-related task demands and variable levels of difficulty is required.

In this study, we examined SZ and healthy control (HC) participants by adapting a previously established paradigm that required active auditory and audiovisual speech processing under different levels of auditory noise (Schepers et al., 2013). Noise reduces the stimulus signal strength, which interferes with auditory and audiovisual speech recognition performance (Ross et al., 2007a). In contrast to a previous study in SZ, in which auditory and audiovisual words were overlayed by auditory background noise (Ross et al., 2007b), we added noise directly to the auditory stimuli. This is an important advantage because the noise in our study did not serve as a temporal cue for the onset of auditory speech. Taken together, our setup with variable levels of auditory noise that is directly added to the stimuli is well-suited to examine the effects of early neural processing on auditory and audiovisual speech recognition in SZ.

## 2 Materials and Methods

### 2.1 Sample and Clinical data

A sample of 31 patients diagnosed with SZ, as defined by the diagnostic and statistical manual of mental disorders, fifth edition (DSM-5), were recruited at the outpatient units of the Charité – Universitätsmedizin Berlin. Of these, 7 had to be excluded from the final analysis (N=5 for task performance problems, and N=2 for EEG signal related problems, outlined below), leaving 24 patients with SZ. To minimize confounding effects of medication on the EEG (Aiyer et al., 2016), patients taking the following medications were not included in the study: Benzodiazepines, lithium, valproic acid, SSRIs, and haloperidol. A total of 27 HC participants were recruited from the general population, of which 6 were excluded from the final analysis (N=2 for task performance problems, and N=4 EEG signal related problems, outlined below). During recruitment, the two groups were matched for handedness (Oldfield, 1971), education (measured in years), smoking (Fagerström, 2012), age and gender (**Table 1**). All participants were screened for comorbid psychopathology with the German version of the Structured Clinical Interview for DSM-4-TR Non-Patient Edition. For the SZ group, symptom severity at the time of measurement was assessed with the Positive and Negative Symptom Scale (PANSS; Kay et al., 1987). Items from this were grouped according to a five-factor model (Wallwork et al., 2012). It is established that cognitive capacity, including executive function, working memory and motor function are generally reduced in SZ (Dickinson et al., 2008; Sheffield et al., 2018). This has been linked to perceptual deficits (Haenschel et al., 2007), and could thus be an instrumental factor affecting performance in the perceptual tasks. Therefore, it is important to have a psychometrically detailed picture of cognitive deficits. In this study, the participant’s cognitive capacity was measured with the Brief Assessment of Cognition in Schizophrenia (BACS; Keefe et al., 2004). Written informed consent was obtained from all participants, including information regarding European Union data protection laws. All participants had normal hearing and normal/corrected to normal vision. People with neurological disorders or head injury with loss of consciousness were excluded at the early recruitment stages, as were participants in the HC group who reported immediate family members with psychiatric or neurological disorders. Drug screening (Drug-Screen Multi 5 Test, Nal von Minden, amphetamine, benzodiazepine, cocaine, opioids, and cannabis) was carried out for all participants and all screenings were negative. The study was conducted in accordance with the 2008 Declaration of Helsinki and approved by the ethics commission of the Charité – Universitätsmedizin Berlin (Approval number: EA1/169/11).

**Table 1:**
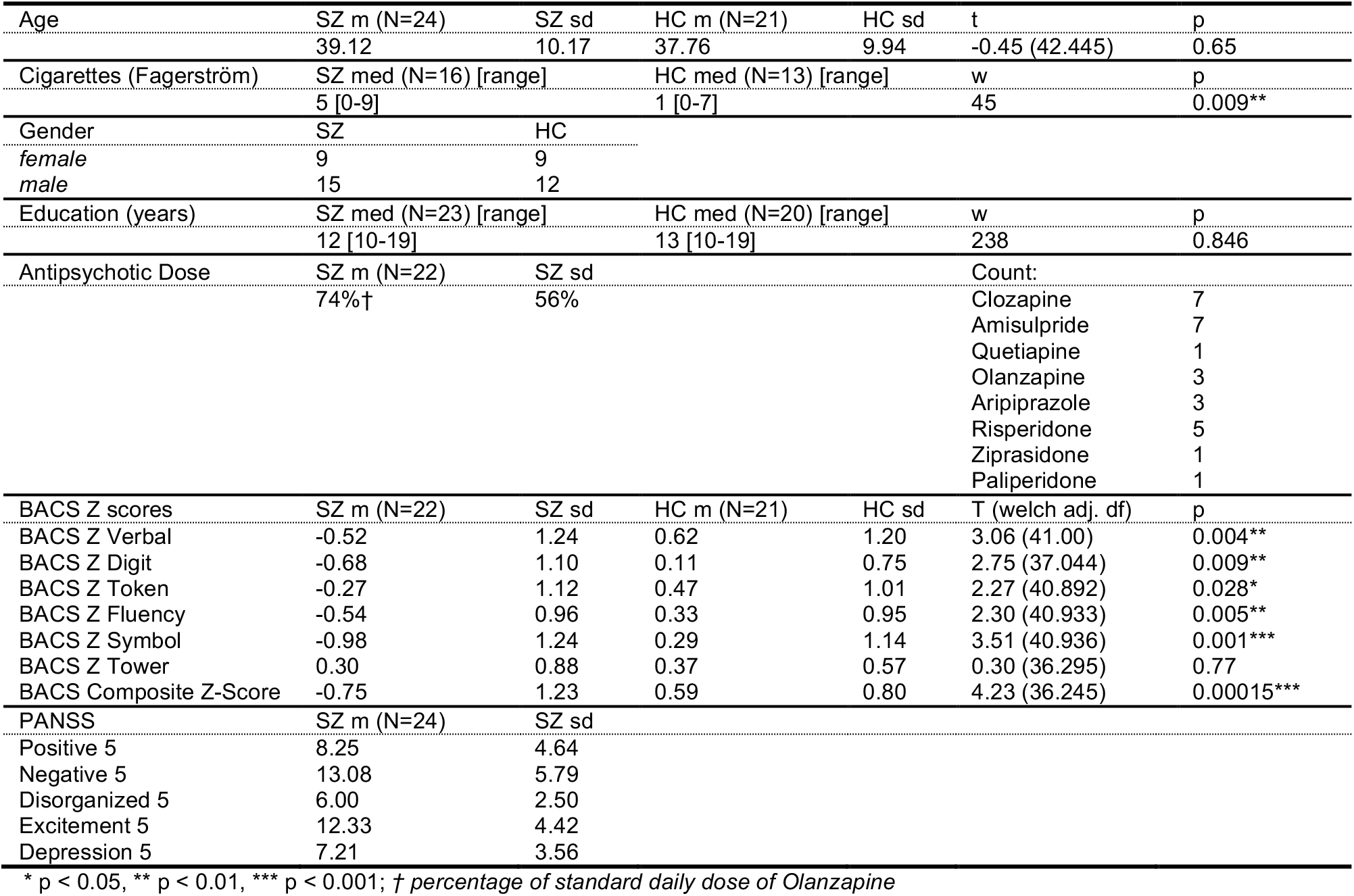
Demographics, medication status, neuropsychological assessment and symptomatology of study participants. Group differences were calculated by either parametric independent t-tests (t) or non-parametric Wilcoxon tests (w), where assumptions for parametric tests were violated. Antipsychotic dose is calculated as the percentage of a standard daily dose of Olanzapine, which is 10mg. BACS raw scores were converted to z-scores, normalized according to age and gender. m = mean, med = median, sd = standard deviation.

|  |  |  |  |  |  |  |
| --- | --- | --- | --- | --- | --- | --- |
| Age | SZ m (N=24) | SZ sd | HC m (N=21) | HC sd | t | p |
|  | 39.12 | 10.17 | 37.76 | 9.94 | -0.45 (42.445) | 0.65 |
| Cigarettes (Fagerström) | SZ med (N=16) [range] |  | HC med (N=13) [range] |  | w | p |
|  | 5 [0-9] |  | 1 [0-7] |  | 45 | 0.009** |
| Gender | SZ | HC |  |  |  |  |
| female | 9 | 9 |  |  |  |  |
| male | 15 | 12 |  |  |  |  |
| Education (years) | SZ med (N=23) [range] |  | HC med (N=20) [range] |  | w | p |
|  | 12 [10-19] |  | 13 [10-19] |  | 238 | 0.846 |
| Antipsychotic Dose | SZ m (N=22) | SZ sd |  |  | Count: |  |
|  | 74%† | 56% |  |  | Clozapine | 7 |
|  |  |  |  |  | Amisulpride | 7 |
|  |  |  |  |  | Quetiapine | 1 |
|  |  |  |  |  | Olanzapine | 3 |
|  |  |  |  |  | Aripiprazole | 3 |
|  |  |  |  |  | Risperidone | 5 |
|  |  |  |  |  | Ziprasidone | 1 |
|  |  |  |  |  | Paliperidone | 1 |
| BACS Z scores | SZ m (N=22) | SZ sd | HC m (N=21) | HC sd | T (welch adj. df) | p |
| BACS Z Verbal | -0.52 | 1.24 | 0.62 | 1.20 | 3.06 (41.00) | 0.004** |
| BACS Z Digit | -0.68 | 1.10 | 0.11 | 0.75 | 2.75 (37.044) | 0.009** |
| BACS Z Token | -0.27 | 1.12 | 0.47 | 1.01 | 2.27 (40.892) | 0.028* |
| BACS Z Fluency | -0.54 | 0.96 | 0.33 | 0.95 | 2.30 (40.933) | 0.005** |
| BACS Z Symbol | -0.98 | 1.24 | 0.29 | 1.14 | 3.51 (40.936) | 0.001*** |
| BACS Z Tower | 0.30 | 0.88 | 0.37 | 0.57 | 0.30 (36.295) | 0.77 |
| BACS Composite Z-Score | -0.75 | 1.23 | 0.59 | 0.80 | 4.23 (36.245) | 0.00015*** |
| PANSS | SZ m (N=24) | SZ sd |  |  |  |  |
| Positive 5 | 8.25 | 4.64 |  |  |  |  |
| Negative 5 | 13.08 | 5.79 |  |  |  |  |
| Disorganized 5 | 6.00 | 2.50 |  |  |  |  |
| Excitement 5 | 12.33 | 4.42 |  |  |  |  |
| Depression 5 | 7.21 | 3.56 |  |  |  |  |

\* p < 0.05, \*\* p < 0.01, \*\*\* p < 0.001; † percentage of standard daily dose of Olanzapine

### 2.2 Setup and Procedure

Participants sat in an electrically and acoustically shielded chamber with low lighting. The experiment was programmed in PsychToolBox (Brainard, 1997; Pelli, 1997) and there were six experimental conditions: Three unisensory auditory (A) conditions (no, low, and high noise) and three bisensory audiovisual (AV) conditions (no, low, and high noise; see Figure 1). Three syllables were used as stimuli: /da/, /ga/, and /ta/. At the beginning of each block, the participants were instructed to respond to a particular target syllable. Within each block, the target probability was lower than the probability of the other two standard syllables (∼23% and 38.5%, respectively). This ensured a larger number of standards for the final analysis, with ∼160 trials for each of the 6 conditions (noise: no, low, high; mode: AV, A). The visual clips had a total duration of 600ms and they were made of 17 different frames with a duration of ∼35ms, (CRT monitor refresh rate was 85 Hz).

**Figure 1:**
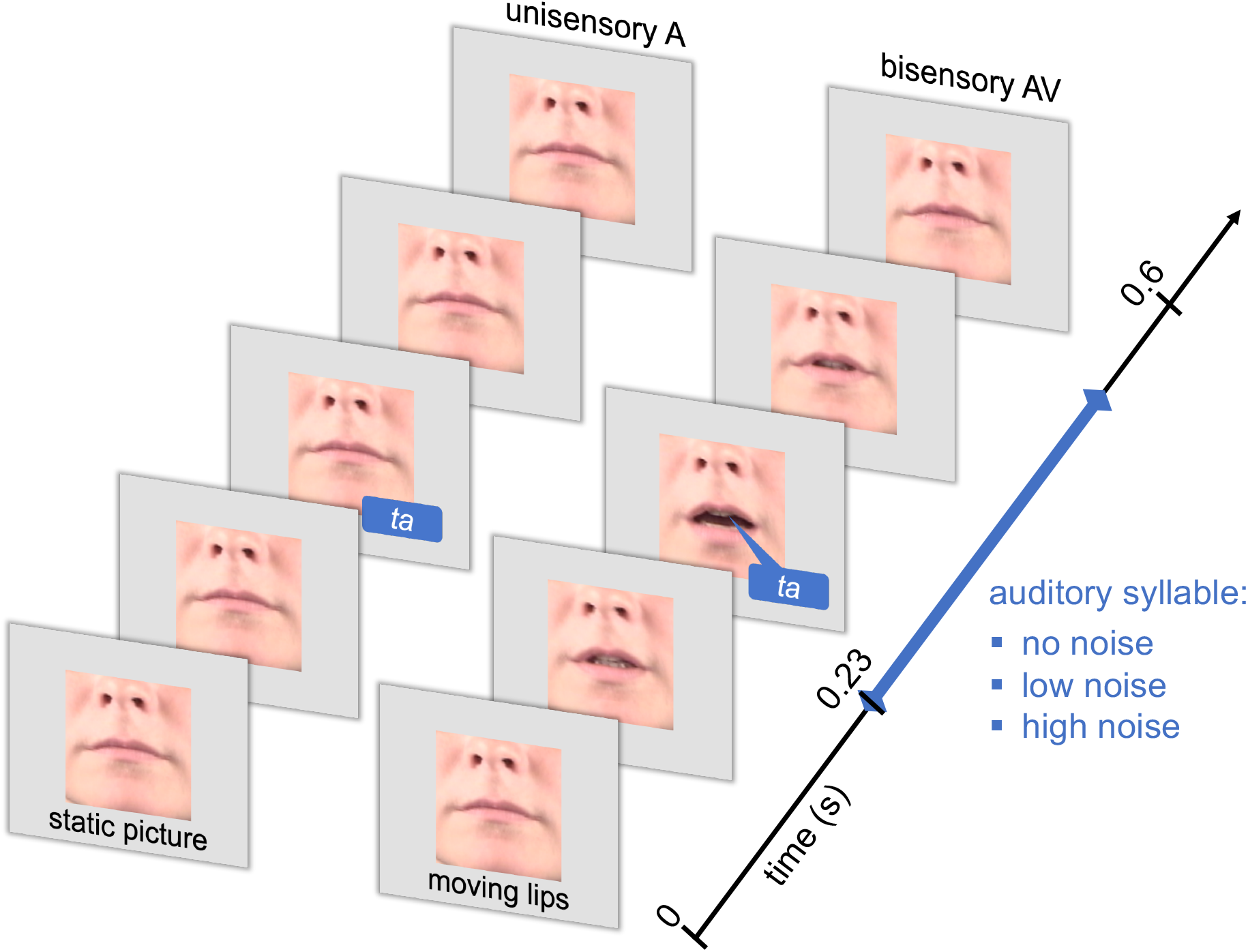
Illustration of a unisensory A and a bisensory AV trial. In unisensory A trials (upper panel) a static picture is presented throughout the trial. In bisensory AV trials (lower panel) visual motion commenced 230ms before auditory stimulus onset. Auditory stimuli, which were presented at three noise levels (no, low, high) consisted of three different syllables (/da/, /ga/, /ta/). Participants were required to identify one of these in each block.

The three syllables were voiced by a female speaker. The signal had a length of 270 to 290ms (sampled at 44.1 kHz). The program ‘Cool Edit Pro (Adobe Systems)’ was used to edit the sound files. The syllables were presented over in-ear air pressure headphones (66dB, E-A-R tone Gold, Auditory Systems, Indianapolis, USA). For the conditions involving auditory noise, the syllable sounds were degraded with additional noise. This was generated by estimating the temporal power signal as the smoothed, rectified auditory signal (convolution with a Hanning window of a temporal width of 5.7ms) and spectral power distribution as the smoothed power spectrum (convolution with a Hanning window of spectral width of 630 Hz). Each syllable was edited separately. Following this, syllable-specific noise with matching spectral and temporal power distributions were generated. In a final step, stimuli with different levels of degradation were generated by altering noise and audio signal with different weights whilst maintaining a constant power level. To minimize the chance that a certain pattern of noise would become recognizable by participants, ten different realizations of the noise were randomly presented for each of the two different noise levels. Since the different syllables are differentially affected by noise levels, we altered the relative weight of the noise for each syllable based on our earlier experiment (Schepers et al., 2013) as follows: /da/ low noise: M = 0.54, /da/ high noise: M = 0.62; /ga/ low noise: M = 0.58, high noise: M = 0.66, and /ta/ low noise: M = 0.66, high noise: M = 0.78. In contrast to our earlier experiment, in which we individually adapted the noise levels, we kept the auditory noise levels with the different noise conditions constant for all participants. This was important because it allows a direct comparison of behavioral and EEG data between groups.

The visual stimuli consisted of short video clips of a female speaker, showing the mouth and nose. The size was 4×4 degrees visual angle, presented against a grey background. The viewer was approximately 120cm distant from the monitor. The images generated were averaged for each frame separately over the three syllables (/da/, ta/, /ga/). Thus, the same visual input was provided for each of the three syllables and it contained no syllable-specific information. In bisensory AV trials the onset of the lip movements always preceded the sound onset by 230 ms. In unisensory A trials a static picture of a nose and a closed mouth was presented instead of a video clip (Figure 1). There was a pause of 0-400ms (mean 200ms) on the first frame and a 1400ms pause on the last frame to allow for responses. Thus, each trial was on average 2200ms. Varying the time interval of the initial pause was important to ensure that the onset of the first frame did not serve as a temporal cue for the onset of the auditory syllable. There were a total of 1248 trials presented, divided into a total of twelve blocks (104 trials per block) of about 3 minutes and 48 seconds each.

### 2.3 EEG Recordings

Data were recorded with a 128-channel passive EEG system (EasyCap). This included two electrooculography (EOG) electrodes (online: 1000 Hz sampling rate with a 0.016-250 Hz bandpass filter; offline. 49-51 Hz, 4^th^ order Butterworth notch filter, 125 Hz 24^th^ order finite impulse response (FIR) lowpass filter, downsampled to 500 Hz, .3 Hz 1500^th^ order FIR highpass filter). The offline nasal reference was re-referenced online to the average of all the EEG electrodes. Nonstationary artifacts were identified and cut with visual inspection. There was no difference between HC and SZ groups in number of remaining trials (SZ: M = 1140.54, [SD = 117.58], HC: M = 1118.05, [SD = 109.76], t(43.80) = −0.66, p = 0.511). Independent component (IC) analysis was used to remove eye movement artifacts as well as heart rate artifacts (Chaumon et al., 2015; Lee et al., 1999). There was no difference between groups in number of components removed (SZ: M = 4.25, [SD = 1.98], HC: M = 4.24, [SD = 0.94], t(33.83) = −0.03, p = 0.979). Noisy channels were cut and then interpolated with spherical interpolation, there were no differences between groups (SZ: M = 10.92, [SD = 4.49], HC: M = 11.24, [SD = 4.19], t(42.80) = 0.25, p = 0.805). However, during preprocessing 6 participants (SZ:2, HC: 4) had to be removed from further data analysis due to excessive eye-blinks or sweat artifacts.

### 2.4 Analysis of Behavioral Data

The analysis of behavioral data focused on d-prime values and hit-rates. The d-prime values for behavioral responses were calculated from hit-rates and false-alarm rates. Hit rates were defined as the proportion of correct responses to the relevant target within each of the 6 conditions (3 noise levels by 2 mode levels). The false-alarm rate was the proportion of false responses under a given target condition within each of the 6 conditions. Behavioral responses outside of the range of 100-900ms following the onset of the auditory syllable were removed from the d-prime calculation. Across groups, the probability of a response outside of this time range was relatively low (SZ: Med = 1.74 % [range: 0.34-7.64 %] and HC: Med = 0.69 % [range: 0-4.17 %], W = 133.5, p = 0.018). Seven participants were cut owing to performance problems (HC = 2, SZ = 5). In these participants the d-primes were uniformly at chance levels owing to either difficulties understanding the task or technical problems with the online registration of stimulus triggers. The d-prime values and hit-rates were analyzed separately with linear mixed effects models (Bates et al., 2015), specifically using the lmerTest package (Kuznetsova et al., 2017) in R (3.6.3), which offers approximations of p-values and degrees of freedom via Satterthwaite’s method. For both models, the independent factors Group (SZ vs. HC), Mode (AV vs. A), and Noise (no, low, high) were entered as fixed variables, with all two- and three-way interactions. As random effects we made intercepts for individual participants. Follow-up Bonferroni-adjusted contrasts were calculated with the emmeans package.

### 2.5 Analysis of Evoked Brain Activity

For the calculation of ERPs, high-pass filtering (2 Hz, 4^th^ order Butterworth) and low pass filtering (45 Hz, 14^th^ order Butterworth) was applied offline to remove low frequency drifts and better define our components of interest, i.e., N1 and P2. The baseline was set at between − 430ms and −330ms relative to auditory stimulus onset. This baseline was necessary to avoid overlapping with visual movement onset in the AV conditions, which began at −230ms relative to auditory stimulus onset. Based on previous studies examining audiovisual speech processing in ERPs (Baart et al., 2014; Schepers et al., 2013; Shahin et al., 2012; van Wassenhove et al., 2005), an ad-hoc defined mediofrontal region of interest (ROI), comprising 14 electrodes, was used for the statistical analysis. To define the times of interest (TOIs) all 6 experimental conditions were combined to create group means. Then, a weighted overall mean was calculated that accounts for the unequal sample size between SZ and HC. Figure 2 illustrates the ROI and the weighted grand-average over groups and conditions. For the grand-average ERP the minimum amplitude of the N1 was observed at 100ms and the maximum amplitude of the P2 was found at 198ms post auditory onset. Accordingly, the TOI for the statistical analysis of the N1 component was defined as 85-115ms, and the TOI for the broader P2 component was defined as 178-218ms.

**Figure 2:**
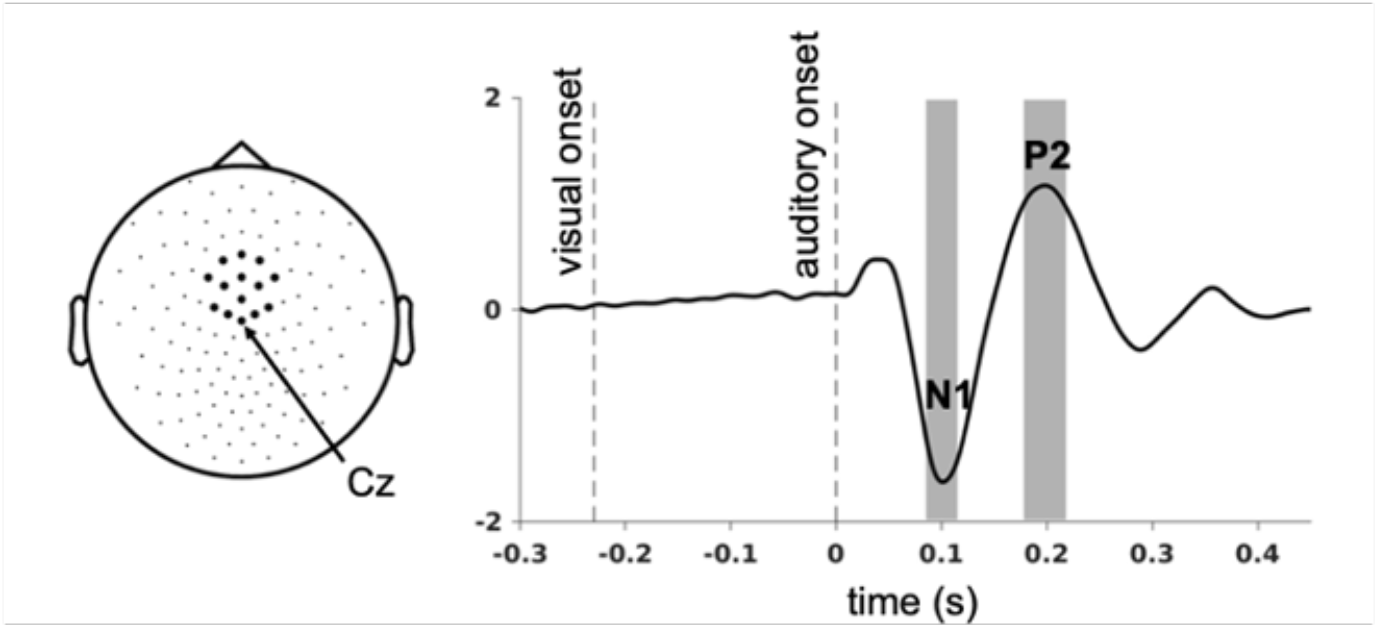
Event-related potentials averaged across groups and experimental conditions at mediofrontal scalp. Left panel: The mediofrontal ROI comprised of 14 neighbouring electrodes, including Cz. Right panel: Weighted grand-average ERP computed across participants (SZs and HCs) and experimental conditions (6 conditions). The N1 and theP2 components time-locked to the onset of auditory syllables are clearly visible at the mediofrontal ROI. The vertical dotted lines indicate the onset of the lip movement in the bisensory AV condition (−230 ms) and the onset of the auditory syllable (0 ms). The vertical gray bars illustrate the time intervals selected for the analysis of the N1 (85-115 ms) and P2 (178-218 ms) components.

LMEs (ImerTest package; Kuznetsova et al., 2017) were applied to the mean amplitudes at these ROIs and TOIs with fixed factors Group (SZ vs. HC), Mode (AV vs. A), and Noise (no, low, high), and all interactions, with random intercepts applied for individual participants. The mean amplitudes at these ROIs and TOIs were extracted and analyzed against independent factors Group (SZ vs. HC), Mode (AV vs. A), and Noise (no, low, high), using the lmer package in R (3.6.3) and follow-up Bonferroni-adjusted contrasts with the emmeans package. Where the interpretation of a null result was critical, Bayes Factors (BF10) were calculated to further specify the relative evidence for H0 and H1 (Aczel et al., 2017; Rouder et al., 2009). BF10s between 1 and 3 indicate anecdotal support for the alternative hypothesis (H1); BF10s from 3 to 10 suggest moderate support; and BF10 above 10 indicates strong support for H1. A BF10 of 1 indicates equal support for H1 and H0, BF10 between 1/3 and 1, 1/10-1/3, and below 1/10 suggest respectively anecdotal, moderate, and strong support for H0.

To more precisely test the relation between behavioral performance and ERPs, LMEs were applied for N1 and P2 amplitudes with dprime, mode and group and their interactions as fixed factors and random effect of participants modelled as individual intercepts. we elected to leave noise out of the analysis in view of the sample size, and focusing on the interactions between multisensory processing and our behavioral parameter of interest. In the same way, potential influence of SZ symptomatology, quantified through PANNS, as well as that of cognitive performance level, via BACS, upon the ERPs was also tested with LMEs.

The amplitudes of event-related potentials (ERPs) were correlated with potential confounding factors (olanzapine equivalent dose, Fagerström test). Critical P-values were adjusted for multiple comparisons with Benjami-Hochberg corrections (Benjamini and Hochberg, 1995).

## 3 Results

### 3.1 Behavioral Data

For behavioral data analysis LMEs were applied with the factors Group, Mode, and Noise. The analysis of d-prime values revealed a main effect of Group (F(1,43) = 4.80, p = 0.034), with HC (adj-M = 2.54, [CI: 1.97, 3.11]) having higher d-prime values than SZ (adj-M = 1.80, [CI: 1.27, 2.33]; t(43) = 2.19, p = 0.034, BF10 = 1.95) (Figure 3A). The main effect of Mode did not reach significance (F(1,215) = 3.44, p = 0.065). Across groups the d-prime values were numerically higher for the AV (adj-M =2.27, [CI: 1.86, 2.67]) compared to the A condition (adj-M = 2.07, [CI: 1.66, 2.48]). There was a main effect of Noise (F(2,215) = 180.10, p < 0.0001), with declining performance with increasing noise level. Importantly, there was also an interaction between Noise and Group (F(2, 215) = 8.01, p = 0.0004). Follow-up contrasts showed that d-primes were higher in HC than SZ, specifically in the no noise condition (HC: adj-M = 4.16 [CI: 3.55-4.78] vs. SZ: adj-M = 2.89 [CI: 2.31-3.46]; t(61.5) = 3.47, p = 0.001, BF10 = 26.87) but not for the low noise condition (HC: adj-m = 2.22 [CI: 1.61-2.84] vs. SZ: adj-m = 1.53 [CI: 0.95-2.10]; t(61.5) = 1.90, p = 0.063, BF10 = 1.23) or the high noise condition (HC: adj-M = 1.22 [CI: 0.61-1.84] vs. SZ: adj-M = 0.99 [CI: 0.41-1.57]; t(61.5) = 0.74, p = 0.527, BF10 = 0.37). No other effects were observed.

**Figure 3:**
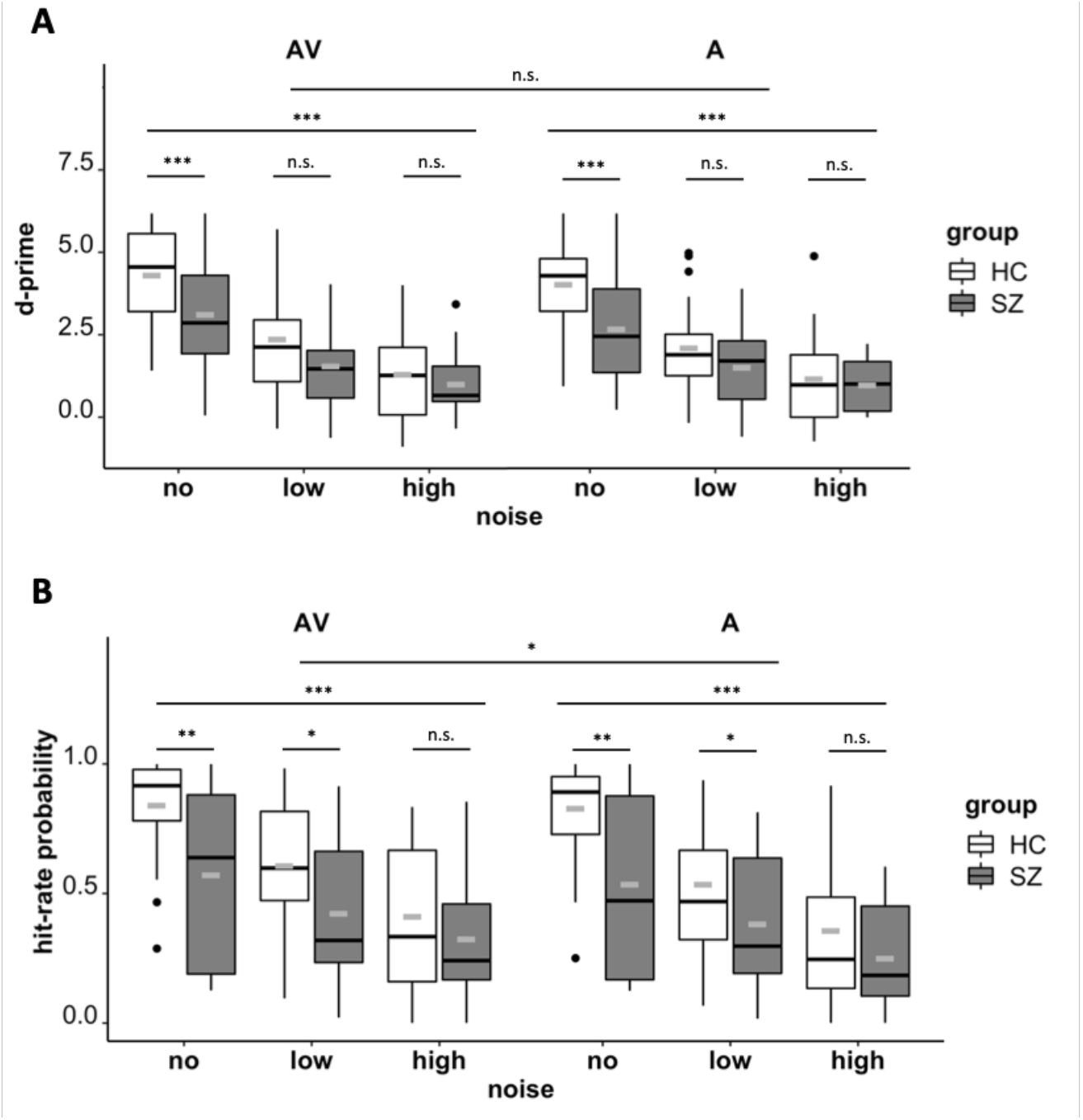
Speech recognition performance differs between bisensory AV vs. unisensory A stimuli, noise levels and study groups. Panel A: d-prime values for bisensory AV (left) and unisensory A (right) stimuli. d-prime values declined with increasing noise levels. Moreover, healthy controls (HC) showed higher d-prime values than SZ, specifically in the no noise condition. No significant group differences were observed for the low and high noise conditions. Panel B: Hit-rates for bisensory AV (left) and unisensory A (right) stimuli. Hit rates declined with increasing noise levels. In addition, hit-rates were higher for bisensory AV compared with unisensory A stimuli. Finally, the hit-rate was lower in SZ than in HC. This difference was most pronounced in no noise and low noise conditions. Gray bar indicates mean values. n.s. p > 0.05, * p < 0.05, ** p < 0.01, *** p < 0.001.

The analysis of hit rates revealed a significant effect of Group (F(1, 43) = 7.78, p = 0.008), showing that HCs (adj-M = 0.60, [CI: 0.49, 0.71]) had higher hit rate probabilities than SZ (adj-M = 0.41, [CI: 0.31, 0.52]) (Figure 3B). There was also a main effect of Mode (F(1, 215) = 5.42, p = 0.021), with higher hit rates in AV (adj-M = 0.53, [CI: 0.45, 0.61) compared to A (adj-M = 0.48, [CI: 0.40, 0.56]) conditions. Moreover, there was a main effect of Noise (F(2, 215) = 100.06, p < 0.0001), with declining performance with increasing noise level. Importantly, the Noise level interacted with Group (F(1, 215) = 6.58, p = 0.002). Follow-up contrasts showed that HC had higher hit rates than SZ in the no noise condition (HC: adj-M = 0.80 [CI: 0.68-0.92] vs. SZ: adj-M = 0.57 [CI: 0.45, 0.69]; t(59.6) = 3.07, p = 0.003, BF10 = 10.73) and the low noise condition (HC: adj-M = 0.57 [CI: 0.44, 0.69] vs. SZ: adj-M = 0.41 [CI: 0.29, 0.53]; t(59.6) = 2.14, p = 0.037, BF10 = 1.79), but not in the high noise condition (HC: adj-M = 0.38 [CI: 0.26, 0.51], SZ: adj-M = 0.29 [CI: 0.17, 0.41], t(59.6) = 1.24, p = 0.221, BF10 = 0.55). No other effects were found.

### 3.2 Evoked Brain Activity

An LME examined N1 amplitudes against factors Mode, Noise, and Group. There was a main effect of Mode (F(1,215) = 40.95, p < 0.0001), with bisensory AV stimuli (adj-M = −1.22 [CI: −1.42, −1.01]) showing less negative amplitude than unisensory A stimuli (adj-M = −1.48, [CI: −1.69, −1.28], t(215) = 6.40, p < 0.0001, BF10 = 107995) (Figures 4-5). This replicates the N1 suppression effect for audiovisual compared to auditory only speech stimuli (Jaaskelainen et al., 2004; van Wassenhove et al., 2005). There was also a main effect of Noise (F(2,215) = 10.99, p < 0.0001), with a decline in N1 amplitude with increasing noise level (Figure 6A). No main effect of Group was found (F(1,43) = 2.33, p = 0.134), but interestingly, there was an interaction between Group and Noise (F(1,215) = 3.46, p = 0.033). Follow-up contrasts revealed larger N1 amplitudes in HC compared to SZ in the no noise condition (HC: adj-M = −1.69, [CI: −2.00, −1.38] vs. SZ: adj-M = −1.28, [CI: −1.57, −0.99]; t(215) = −2.24, p = 0.030, BF10 = 2.12). In contrast, no group differences were found in the low noise condition (HC: adj-M = −1.43, [CI: −1.74, −1.12] vs. SZ: adj-M = −1.19, [CI: −1.47, −0.90]; t(215) = −1.33, p = 0.19, BF10 = 0.60) and in the high noise condition (HC: adj-M = −1.33, [CI: −1.64, −1.02] vs. SZ: adj-M = −1.19, [CI: −1.47, −0.90]; t(215) = −0.77, p = 0.44, BF10 = 0.38). Thus, comparable to the behavioral data, group differences in N1 amplitudes were primarily observed for the no noise condition.

**Figure 4:**
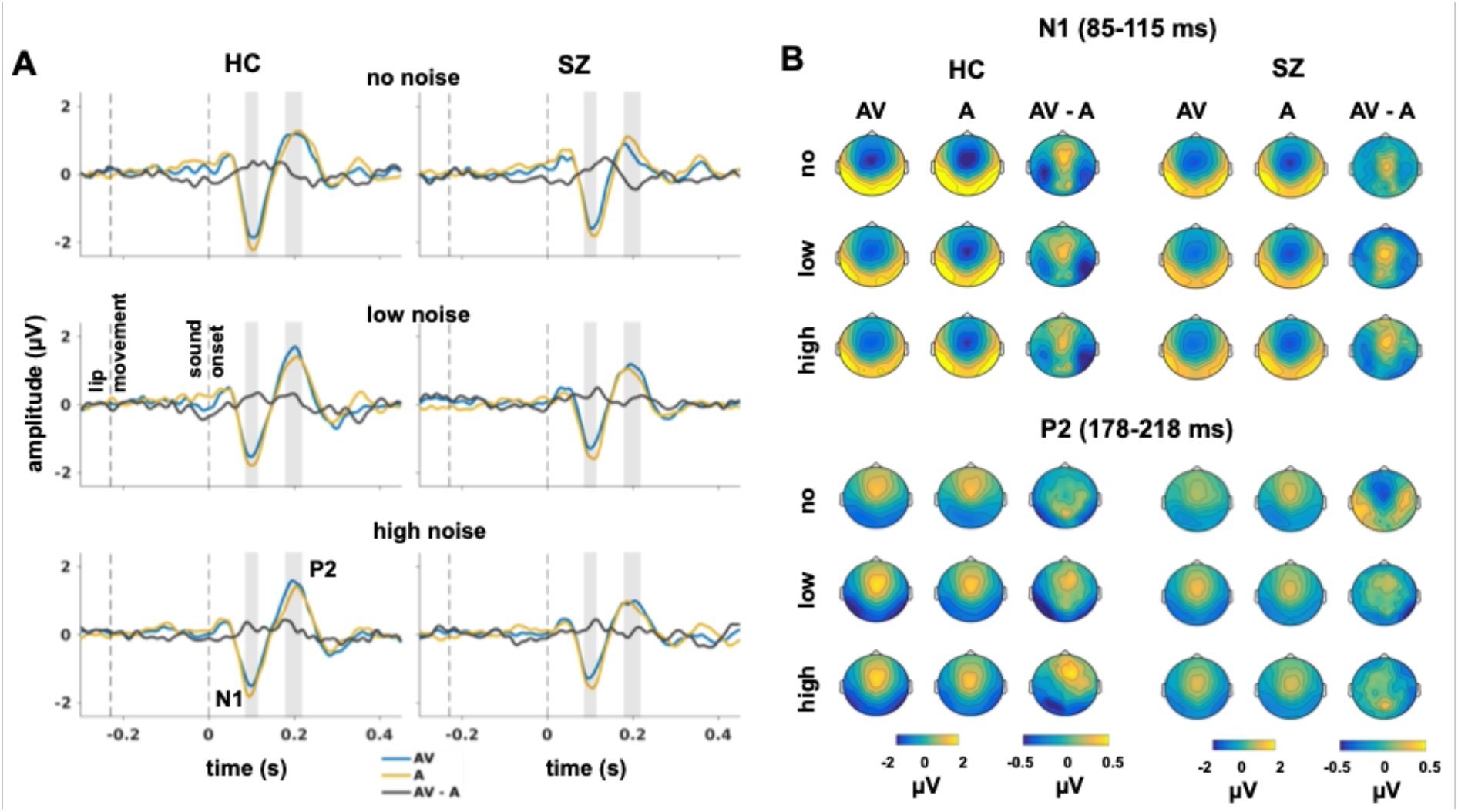
Traces and topographic maps of N1 and P2 components for bisensory AV and unisensory A stimuli. Panel A: Grand average ERP traces of the mediofrontal ROI. For the N1 component, significant smaller amplitudes were observed for bisensory AV compared with unisensory A stimuli. This crossmodal N1 suppression effect was comparable between conditions and groups. Significant group differences in N1 amplitudes were specifically found in the no noise condition, with smaller amplitudes in SZs compared to HCs. No significant group differences were observed in the low and high noise conditions. For the P2 component, reduced amplitudes were found in SZs compared to HC, specifically in response to bisensory AV stimuli, but not in response to unisensory A stimuli. Panel B: Topographic maps of the N1 (upper panel) and the P2 components (lower panel). The plots highlight the mediofrontal topographies of both components.

Another LME tested P2 amplitudes against factors Mode, Noise, and Group. The analysis revealed a main effect of Group (F(1,43) = 4.89, p = 0.032), showing larger P2 amplitudes in HCs compared to SZs (Figure 5B). There were was also main effect of Noise (F(2, 215) = 7.58, p = 0.0007). Low noise (adj-M = 1.10) had larger deflections than no noise (adj-M = 0.90, t(215) = 3.89, p = 0.0004, BF10 = 76.25). High noise (adj-M = 1.01) showed intermediate P2 deflections, which did not significantly differ from the other two other conditions (p > 0.119). Furthermore, there was a main effect of Mode (F(1,215) = 4.00, p = 0.047), with bisensory AV stimuli (adj-M = 1.04), showing larger P2 amplitudes than unisensory A stimuli (adj-M = 0.96, t(215) = 2.00, p = 0.047, BF10 = 1.43). In addition, there was an interaction of Mode by Group, which showed that the effect of Mode was specific to the HC group. The P2 amplitudes in response to bisensory AV stimuli differed significantly between groups (HC: adj-M = 1.26, [CI: 0.99, 1.52]; SZ: adj-M = 0.83, [CI: 0.58, 1.08], t(49.5) = 2.67, p = 0.010, BF10 = 4.66). No Group differences, however, were found for the unisensory A stimuli (HC: adj-M = 1.09, [CI: 0.82, 1.35], SZ: adj-M = 0.83, [CI: 0.58, 1.08], t(49.5) = 1.59, p = 0.118, BF10 = 0.81). There was also an interaction between Noise and Mode. P2 amplitudes in response to bisensory AV stimuli were smaller in the no noise condition (adj-M = 0.87, [CI: 0.65, 1.08]) compared to the low noise (adj-M = 1.16, [CI: 0.94, 1.38], t(215) = −4.13, p = 0.0002, BF10 = 142.81) and the high noise condition (adj-M = 1.10, [CI: 0.88, 1.32], t(215) = −3.29, p = 0.0035, BF10 = 17.61). There was no difference between P2 amplitudes to bisensory stimuli of the low and high noise condition (t(215) = 0.88, p = 1.00). Moreover, no P2 amplitude differences in response to unisensory A stimuli were observed between the different noise levels (all p-values > 0.248). Hence, group differences in P2 amplitudes were specifically observed for bisensory AV stimuli, but did not depend on the noise level.

**Figure 5:**
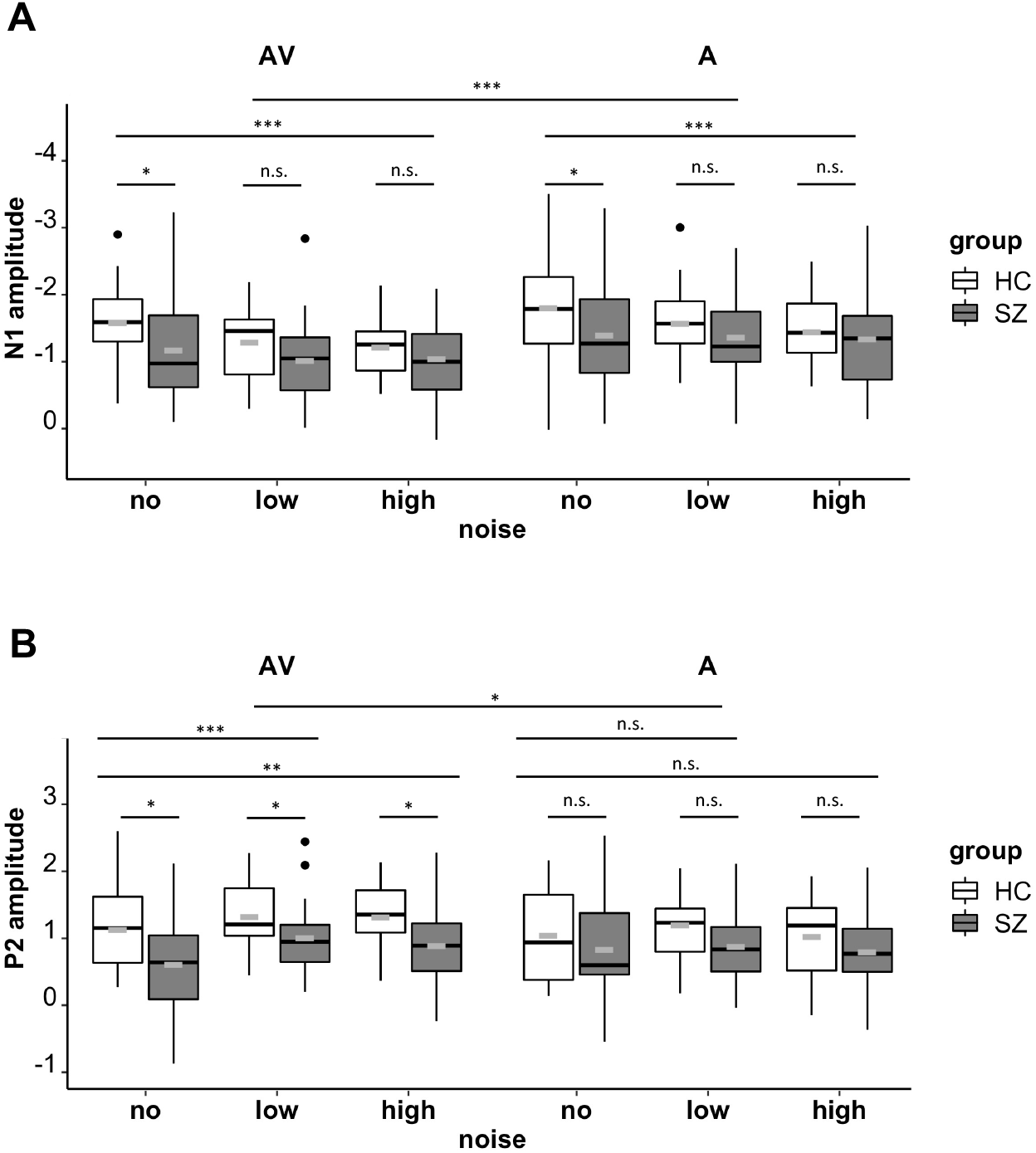
Amplitudes of the N1 and P2 components in response to bisensory AV and unisensory A stimuli. Panel A: Amplitudes of the N1 component. HCs showed higher N1 amplitudes than SZ, specifically in the no noise condition. Across groups and conditions, there was also an N1 suppression effect, with smaller amplitudes for bisensory AV compared to unisensory A stimuli. Moreover, N1 amplitudes declined with increasing noise level. Panel B: Amplitudes of the P2 component. HC showed significantly higher amplitudes than SZ, specifically in the AV condition. Amplitude increased with noise levels specifically in the AV condition. Gray bar indicates mean values. n.s. p > 0.05, * p < 0.05, ** p < 0.01, *** p < 0.001.

LMEs of dprime against N1 showed that overall greater deflections in N1 were associated with overall better dprime performance (F(1, 237.30) = 7.20, p = 0.008). A significant group*dprime interaction (F(1, 237.30) = 5.72, p = 0.0176) suggested that the N1 association was specific to the HC group, with the SZ group showing an undifferentiated response (Figure 6). Follow-up LME analyses for groups separately supported this, showing that higher dprime values were associated with greater deflections for HC (F(1, 110.18) = 18.65, p < 0.0001)), but not for SZ (F(1,126.89) = 0.044, p = 0.835). An LME of P2 on dprime together with group and mode factors showed an overall association of higher dprimes with greater deflections (F(1, 243.30) = 5.98, p = 0.015). No interactions were significant.

**Figure 6:**
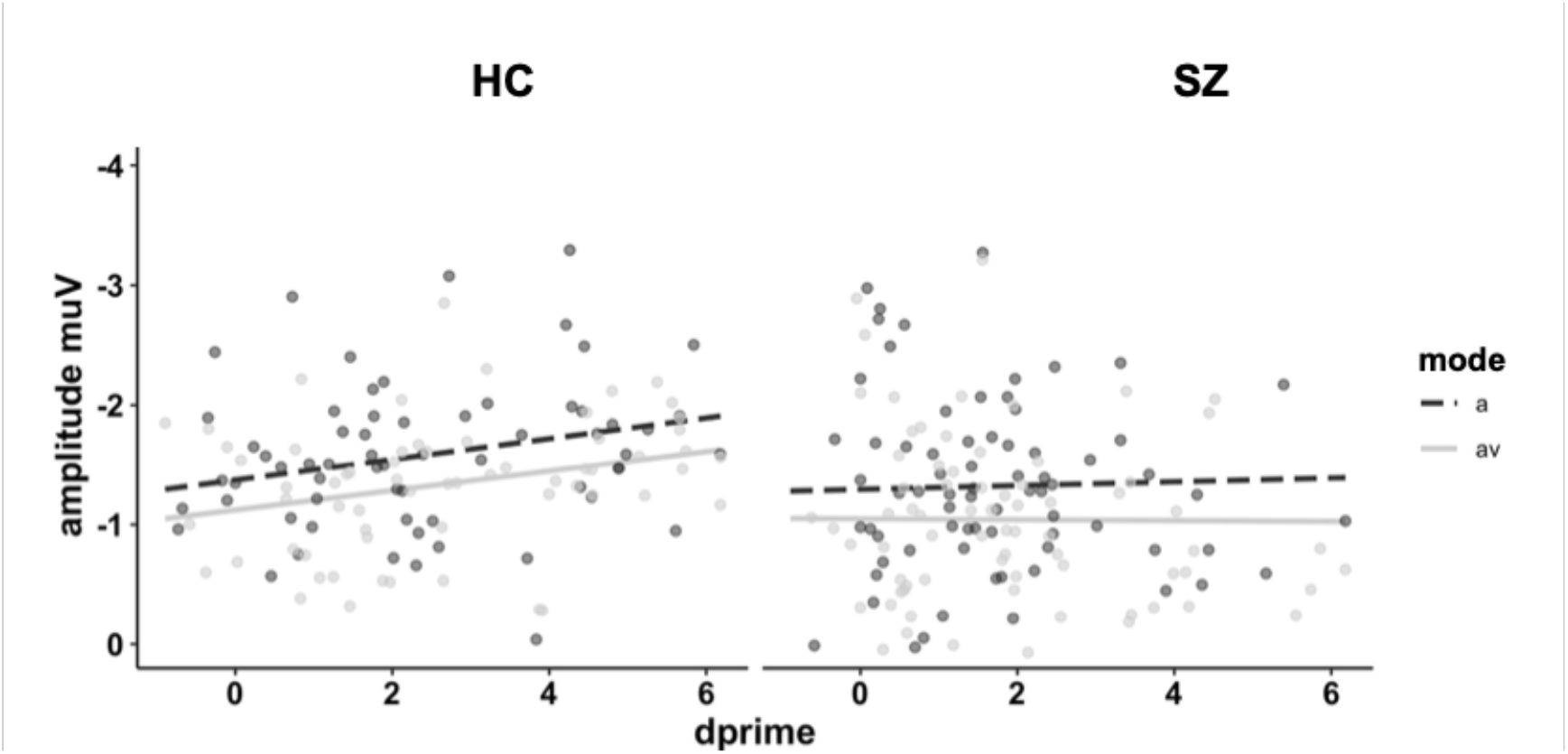
Interaction plot of d-prime against N1 amplitudes across mode and group. The overall main effect suggests that higher behavioral performance scores are associated with greater N1 deflections in response to auditory and audiovisual speech stimuli. This effect was specifically observed in the HC group (left), but not the SZ group (right).

A LME of N1 with the independent variables mode, group, and BACS total z-scores, showed a main effect of BACS (F(1,39) = 4.51, p = 0.040, β = −0.37). This shows that greater cognitive capacity was related to generally higher N1 amplitude deflections (Figure 7). There were no main effects of group (F(1,39) = 0.28, p = 0.599, β = −0.09) or mode (F(1,39) = 3.27, p = 0.078, β = −0.09) and no significant interactions. Two separate LMEs of N1 and also P2 on the five dimensions of the PANSS did not show any significant relations (p > 0.064). Finally, potential confounds psychopharmaceutical medication (olanzapine equivalent dose) and smoking (Fagerström test) were correlated with ERP amplitudes (N1 and P2), with critical p-values adjusted for multiple comparisons. There were no significant correlations between medication and neural parameters (adjusted p > 0.416, BF10 ranged between 0.47-0.64, with the exception of AV high noise and P2, with BF10 = 2.88 suggesting anecdotal support for H1). There was no relation between smoking and neural parameters (adjusted p > 0.63, most BF10 scores ranged between 0.52-1.03. There were two exceptions in the HC group, suggesting anecdotal support for H1 in P2: AV low BF10 = 2.16, AV high BF10 = 1.79).

**Figure 7:**
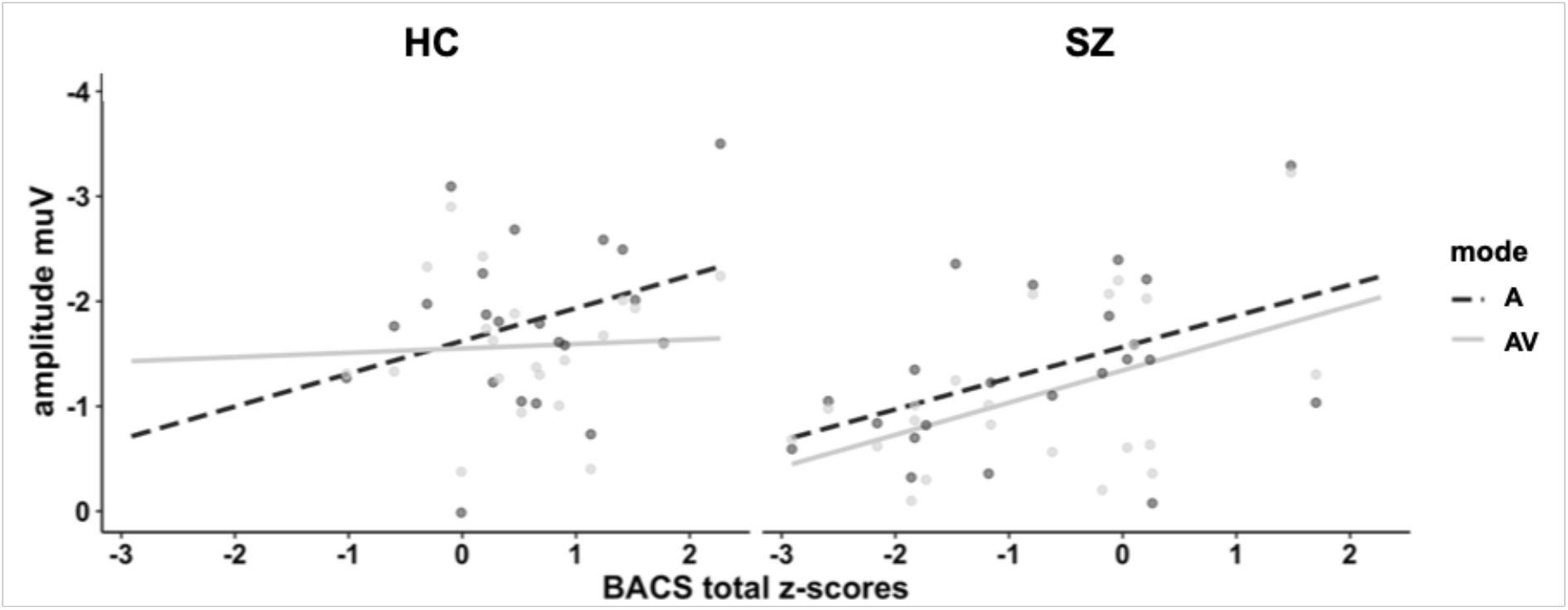
Interaction plot of BACS total z-scores against N1 amplitudes across modality and group. BACS total scores were transformed to z-scores taking into account age and gender. The overall main effect suggests that higher cognitive performance scores are associated with greater N1 deflections in response to bisensory AV and unisensory A stimuli.

## 4 Discussion

In this study we examined early auditory and audiovisual speech processing under different noise levels in HC and SZ. We observed reduced speech recognition performance in SZs in the no noise condition, while the performance differences were diminished or absent in the low and high noise conditions, respectively. Mirroring the behavioral results, we observed reduced N1 amplitudes to auditory and audiovisual speech stimuli in the SZ group, specifically in the no noise condition. In the HC group the N1 amplitudes were related to the behavioral performance, whereas no such relationship was found in the SZ group. In relation to multisensory processing, we observed similar N1 suppression effects for bisensory compared to unisensory speech stimuli in SZs and HCs. Finally, we observed generally positive relationships between the N1 amplitudes and cognitive performance, as obtained by the BACS, in both study groups.

### 4.1 Noise Influences Auditory and Audiovisual Speech Recognition Deficits in Schizophrenia

As expected, increasing the auditory noise level led to a decline in speech recognition performance. In addition, a better performance was found for bisensory audiovisual compared to unisensory auditory speech stimuli. These observations replicate our previous findings in healthy young adults (Schepers et al., 2013). The effects of sensory stimulation, i.e., bisensory vs. unisensory stimulation, on speech recognition did not significantly differ between groups, which indicates a relatively intact multisensory benefit in SZs (Moran et al., 2021). Across experimental conditions, SZs showed a poorer performance than HCs, which supports the notion of a speech recognition deficit in SZ (DeLisi, 2001).

The most interesting finding was that SZs’ relative deficit in speech recognition compared to HCs, declined with increasing noise level. In SZs, compared to HCs, we observed strong group differences in the no noise condition, small group differences in the low noise condition and no group differences in the high noise condition. Thus far, studies examining the effects of auditory noise on auditory and audiovisual speech recognition are sparse (Ross et al., 2007b). In a behavioral study investigating the effects of auditory background noise on auditory and audiovisual speech recognition, Ross et al., (2007b) observed a diminished multisensory benefit of viewing lip-movements in SZ compared to HC at an intermediate noise level, which fits with the principle of inverse effectiveness (Senkowski et al., 2011; Stein and Meredith, 1993; van de Rijt et al., 2019). Critically, the background noise in this study always started at a fixed interval of 520ms prior to the auditory onset, i.e., around the onset of the lip movements. Thus, the noise in this study served as a reliable temporal cue for the auditory speech onset. Given the evidence that temporal multisensory processing is compromised in SZ (Stevenson et al., 2017), it is possible that group differences in temporal processing influenced the findings by Ross et al. (2007b). In our study, the noise was directly added to the auditory syllable and did therefore not provide temporal information about the auditory speech onset. There are also differences in speech recognition performance between the previous findings and those of our study. In Ross et al. (2007b) the hit rate for no noise stimuli in both study groups was above 90% for bisensory stimuli and around 85% for unisensory auditory stimuli. In our study, the hit-rate for combined bisensory and unisensory stimuli in the no noise condition was on average 80% in HCs and 57% in the SZ group. Thus, in the bisensory noise condition in which SZs showed multisensory gain deficits in Ross et al. (2007b), the hit rate was comparable to the hit rate observed in the no noise condition in our study. This implies comparable processing demands, which could have contributed to the speech recognition deficits in the Ross et al. (2007b) and in the present study. In summary, our study revealed aberrant speech recognition for auditory and audiovisual speech stimuli in SZ, which, however, declined with increasing noise level.

### 4.2 Early Evoked Brain Activity Relates to Speech Recognition Deficits in Schizophrenia

In agreement with previous studies emphasizing the role of early evoked brain activity for auditory and audiovisual speech processing in healthy individuals (Baart, 2016; Baart et al., 2014; Bernstein et al., 2008), our analysis focused on the N1 and P2 components triggered by the onset of the auditory syllables. In parallel with our behavioral results, we observed group differences in N1 amplitudes for both auditory and audiovisual speech stimuli SZ, primarily in the no noise condition. In contrast, group differences in N1 amplitude were attenuated in the low and high noise conditions. Previous studies using non-speech unisensory auditory stimuli (Gallinat et al., 2002; Leicht et al., 2010) as well as speech stimuli (Mathalon et al., 2019; Perez et al., 2012) have shown reduced N1 amplitudes in SZ. Given that alterations in N1 amplitudes have been found in first-episode SZs as well as clinically unaffected first-degree relatives, it has been hypothesized that reduced auditory N1 amplitudes are an endophenotype of SZ (Foxe et al., 2011; Salisbury et al., 2010).

Our observation of reduced N1 amplitudes in the no noise condition in SZs raise questions regarding the neural basis of this finding. The majority of studies reporting reduced N1 amplitudes in SZ have used salient auditory stimuli like sinusoidal tones (Rosburg et al., 2008). As such, our finding of reduced N1 responses to salient no noise auditory and audiovisual speech stimuli fits with these previous studies. Enhanced trial-by-trial fluctuations of neural activity in sensory and higher order cortical areas have been consistently observed in SZ (Javitt and Sweet, 2015; Rolls et al., 2008). Furthermore, it is likely that excitation/inhibition (E/I) imbalance in neural circuits contributes to the enhanced fluctuations in neural activity (Anticevic and Lisman, 2017; Uhlhaas and Singer, 2015), which, in turn, could be a factor contributing to reduced evoked brain responses (Hall et al., 2015; Winterer et al., 2000). It is possible that enhanced trial-by-trial fluctuations in neural activity primarily affected the processing of the salient speech stimuli in SZs. In our study, we found positive relationships between the N1 amplitudes and behavioral performance in the HC group but not in SZs. Hence, it may be that the N1 amplitudes in SZs reflect a stronger bias of non-speech related factors, such as variations in attention or memory-related processing. The correlations with speech recognition performance also indicate a better decoding of the speech signals in the N1 amplitudes, which are generated in the auditory cortex and to some extent in higher order areas (Bernstein et al., 2008; Besle et al., 2008; Campbell, 2008), in HCs compared to SZs. Future research implementing multivariate decoding approaches could provide further insight into this interesting question (Bae et al., 2020). Finally, in both groups we observed positive relationships between the N1 amplitudes and cognitive performance, as obtained by the BACS total score. In a recent study, positive relationships with the BACS total score have been found for the auditory discrimination threshold but not for clinical symptoms in SZs with psychosis (Ramsay et al., 2020). This suggests a link between basic auditory processing, as reflected in the evoked N1 component and auditory discrimination threshold, with general cognitive performance that is not explained by clinical symptoms. Taken together, the findings of reduced N1 amplitudes for no noise auditory and audiovisual stimuli in SZ mirror the behavioral results. This indicates that the early speech processing deficits in SZs primarily relates to salient stimuli.

The findings for the P2 component were more complex than for the N1 component and they were confined to bisensory audiovisual stimuli. In both study groups, the amplitude of the audiovisual P2 component was larger for the no noise compared to the two noise conditions. This observation resembles our previous finding in young healthy adults (Schepers et al., 2013). Moreover, for all three noise conditions, we observed larger audiovisual P2 amplitudes for HCs compared to SZs. Given that no such effects were found for the unisensory stimuli, this indicates group differences in the multisensory influence of the visual inputs on auditory speech processing. Nevertheless, we did not find any relationships between the P2 amplitudes with task performance or cognitive functions. Hence, the observed group differences on the bisensory P2 were relatively unspecific and they were not directly relevant for behavior or cognition.

#### Multisensory Benefits are Comparable in SZs and HCs

Another important aspect of our study is the question regarding multisensory integration differences between SZs and HCs. Reduced N1 and P2 amplitudes for bisensory compared to unisensory stimuli are consistent findings in audiovisual speech research (Baart et al., 2014; Besle et al., 2004; Jaaskelainen et al., 2004; van Wassenhove et al., 2005). Previously, Stekelenburg et al. (2013) examined the N1 suppression effect for no noise audiovisual vs. auditory stimuli in SZs and HCs. In contrast to our study, the authors did not find an N1 suppression effect in SZs. Recently, we observed a close interplay of top-down attention and multisensory integration in SZ (Moran et al., 2021), which fits with the notion that top-down mechanisms can influence basic integrative sensory processing (Senkowski et al., 2005; Talsma et al., 2010, 2007). In line with this notion, we proposed that differences in attentional demands could have contributed to previous seemingly contradictory findings on multisensory integration in SZs. As such, it could be that the differences in findings between Stekelenburg et al. (2013) and our study relate to differences in top-down attention demands.

In our study there are several findings that suggest a relatively intact multisensory benefit in SZs. First, across the different noise levels the multisensory benefit in speech recognition performance was comparable for SZs and HCs. Second, across the different noise levels there were no differences in bisensory N1 amplitude suppression effects between groups. Third, the behavioral deficits and N1 reductions in the no noise condition in SZs were comparable for unisensory and bisensory stimuli. In summary, our study suggests a relatively intact multisensory benefit in SZ. This implies that the observed deficits in audiovisual speech processing mainly relate to aberrant processing of the auditory syllables.

### 4.3 Limitations

We tested for evidence of the impact of nicotine consumption or current medication on N1 and P2 findings. Although no results were significant even with a less conservative Benjamni-Hochberg adjustment for multiple comparisons, Bayes Factor calculations suggested anecdotal support for a relation in some P2 correlations. In particular, between nicotine consumption and the two higher noise conditions in the HC group, and the two higher AV noise conditions and medication in the SZ group. Given the unavoidably high number of comparisons, these findings are likely to be spurious, and are in any case not related to our central findings relating to N1. We found no relation between SZ psychopathology, as measured by the PANSS, with behavioral or EEG parameters. It is possible that any such relations require a larger sample size than that of our experiment, which was primarily designed to uncover differences between groups. Our measurement of the PANSS was prior to the EEG measure, rather than during the measurement, so any potentially interesting relation between, e.g., auditory hallucinations and task performance and EEG, could not be assessed. Future studies could employ the Audio-Visual Abnormalities Questionnaire (AVAQ; Nikitova et al., 2019), to relate experimental findings to this standardized measure of everyday experiences of audiovisual abnormalities in people with SZ. This interesting instrument was not available at the time of data collection. Finally, it is possible that long-term medication effects in SZs contributed to our findings. Thus, as with other experiments examining chronic SZ patients, a replication of findings in a sample of first-episode and ideally unmedicated patients would be desirable.

### 4.4 Conclusion

Our study demonstrates that impaired early neural processing, as reflected in reduced N1 amplitudes, relates to auditory and audiovisual speech processing deficits in SZ. Our findings also suggest that this impairment is confined to salient speech input. In our study we did not find evidence for alterations in integrative multisensory processing in SZs. In fact, bisensory N1 suppression effects and comparably better speech recognition performance for bisensory compared to unisensory stimuli was consistent across both groups. This suggests a relatively intact multisensory benefit in SZs, which indicates that the observed deficits in auditory and audiovisual speech recognition were primarily related to aberrant processing of auditory speech.

## Acknowledgments

This research was funded by a grant to DS from the German Research Foundation (SE1859/4-1). We would like to thank Julian Keil for his help in programming of the experiment and Alex Masurovsky, Teresa Ramme, Lisa Renziehausen, Marianne Bröker, and Joseph Wooldridge for their assistance gathering the data.

